# Research and conservation priorities to protect wildlife from collisions with vehicles

**DOI:** 10.1101/2022.11.01.514670

**Authors:** Pablo Medrano-Vizcaíno, Clara Grilo, Manuela González-Suárez

## Abstract

The rapidly expanding global road network poses threats to wildlife, including direct mortality. Given limited knowledge and resources, strategic allocation is critical. We introduce a method to identify priority areas and taxa to study and protect affected by vehicle collisions using Latin America as a case study. In this region high biodiversity and an expanding road network can result in high impacts from roads, yet emerging research expertise offers opportunities for action. To identify priority targets we combined predicted spatially-explicit roadkill rates for birds and mammals with information about the current road network and species conservation status. Priority areas for conservation (with many species susceptible to roadkill but few or inexistent roads) were largely concentrated in the Amazon; while priority areas for research (unstudied regions with many roads and many species susceptible to roadkill) occur in various areas from Southern Mexico to Chile. Priority taxa for conservation reflected studied, roadkill-susceptible groups (eg, vultures and armadillos), while priority taxa for research were defined as either poorly-studied roadkill-susceptible groups or unstudied groups of conservation concern (eg, cuckoos and shrew opossums). Our approach offers a tool that could be applied to other areas and taxa to facilitate a more strategic allocation of resources in conservation and research in road ecology.

## Introduction

Roads are widespread features in our planet and already fragment some of the world’s last remaining wilderness areas such as the Amazon (Laurance et al., 2014; Meijer et al., 2018). By 2050, an additional 25 million kilometers of new roads will be constructed primarily in Africa, South and East Asia, and Latin America (Laurance et al., 2014). Roads are one of the main anthropogenic causes of wildlife mortality worldwide (Hill et al., 2019), and the primary cause in some regions (Taylor-Brown et al., 2019). For example, an estimated 194 million birds and 29 million mammals are killed in European roads, 340 million birds in the United States, and twelve million birds and five million mammals in Latin America (Grilo et al., 2020; Loss et al., 2014; Medrano-Vizcaíno et al., 2022).

The relentless expansion of roads affects wildlife worldwide, but impacts can vary across different regions and habitats depending on presence of species more exposed to roads. Research shows evidence that roadkill risk varies among taxa due to their distinct morphological and ecological characteristics. For example, higher roadkill rates are found in larger, ground-foraging birds with more diverse diets and also in medium-sized, diurnal, scavenging mammals (Caceres 2011, González-Suárez et al. 2018). Since species occur in different habitats, there are some regions with more roadkill incidence than others (Medrano-Vizcaíno et al., 2022). These generalized links between road mortality, traits and habitats suggest we could use existing information to predict wildlife mortality patterns across large geographical and taxonomic scales and thus, identify priority targets for research, infrastructure design and planning, and conservation management.

The application of prioritization tools to inform future research actions is particularly valuable in areas where risks are expected to be high (i.e., biodiverse areas with expanding road networks) but where empirical estimates of impacts are scarce. Latin America is a highly biodiverse region harboring eight biodiversity hotspots and unique endemism (Myers et al., 2000) as well as ∼3.5 million km of roads (Meijer et al. 2018). Road impacts threaten Latin-American wildlife, but assessments are still rare with few, albeit rising, road ecology studies (Pinto et al., 2020). This makes Latin America an ideal case study to test the proposed approach capitalizing on existing spatially-explicit roadkill rates predicted for bird and mammal species known to suffer mortality from collisions (obtained from Medrano-Vizcaíno et al., (2022)). These data, combined with information on road network and species conservation status offer a way to identify priority areas and taxa for conservation and research.

To identify priorities, we propose that conservation efforts to reduce road impacts should focus on areas where many species susceptible to roadkill and few roads coincide (high vulnerability, low exposure). On the other hand, priority areas for research should be those currently unstudied but with many species susceptible to roadkill and many roads (high vulnerability, high exposure. Fig. 1). Similarly, we define priority taxa for conservation considering taxonomic orders with at least one third of species recorded as roadkill and high predicted roadkill rates. We propose priority taxa for research should be currently understudied groups for which available roadkill rates are high, as well as unstudied groups in which many species are considered to be of conservation concern (Fig 2). This approach can be applied to other regions and different taxonomic or functional groups offering a tool to guide resource allocation in road ecology and conservation.

**Figure 1.**
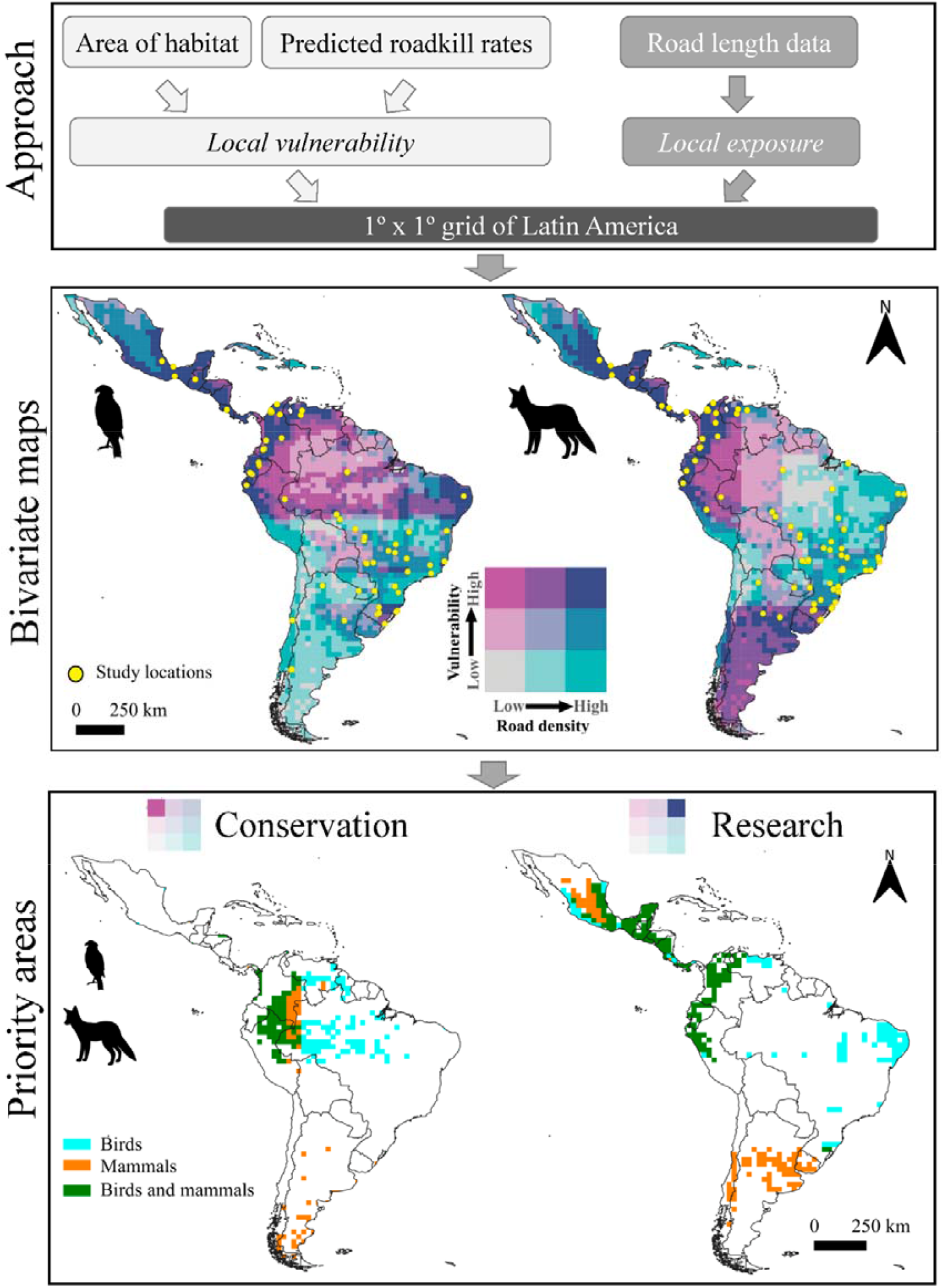
Summary of proposed approach to identify priorities (top panel) with resulting output representing maps of local vulnerability and exposure (middle panel) and the proposed priority areas for conservation and research for Latin American birds and mammals (bottom panel). Middle panel shows bivariate choropleth maps with nine categories based on terciles values for local vulnerability and exposure, with yellow symbols showing where data were collected. Bottom panel shows proposed priority areas for conservation and research based on bivariate maps (the modified legends on top of this panel highlights that we considered high vulnerability and low exposure for conservation priorities, and high vulnerability and high exposure for research priorities). Animal silhouettes were obtained from Phylopic (http://phylopic.org).

**Figure 2.**
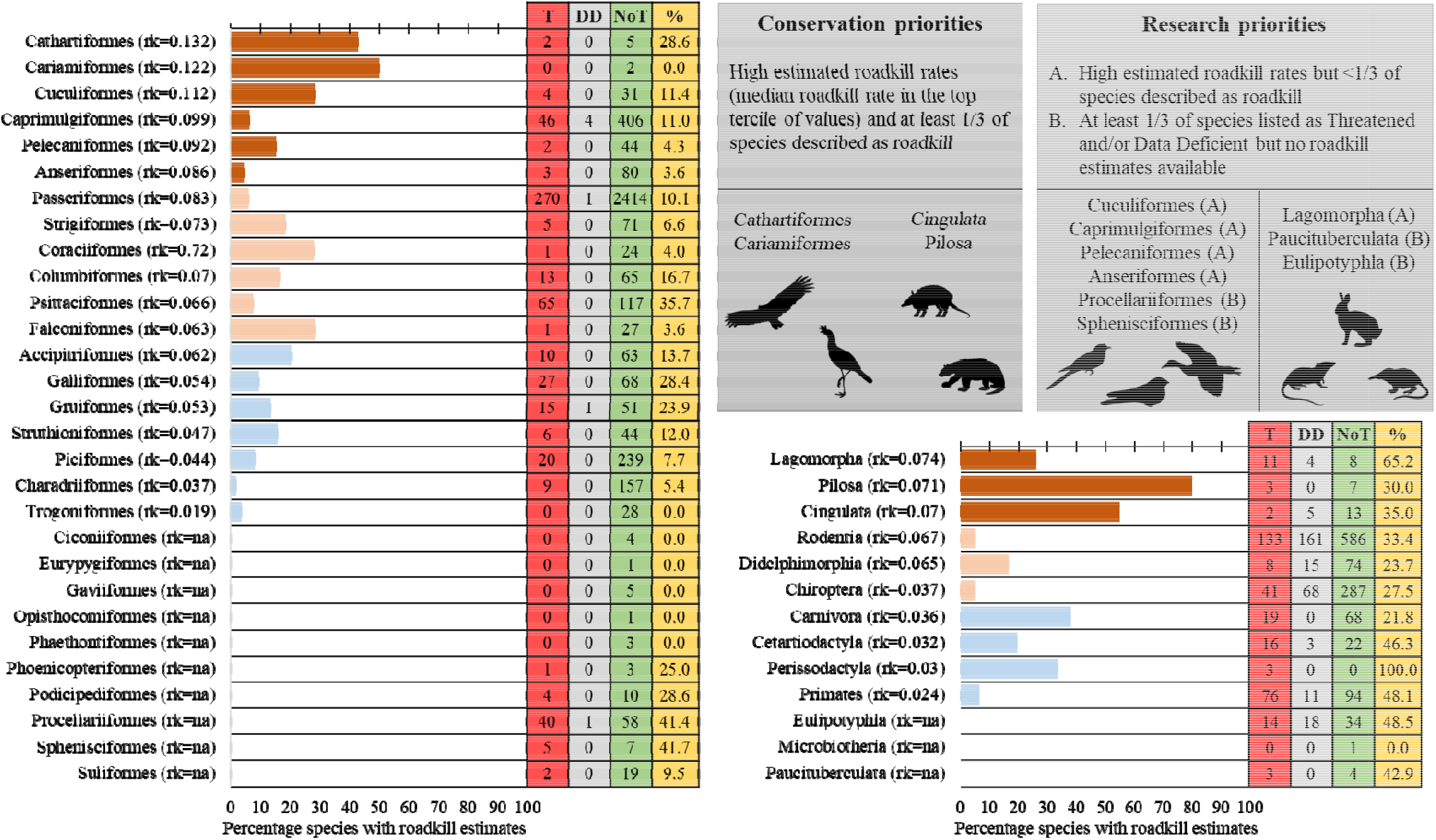
Priority conservation and research taxa and criteria values for bird and mammal taxonomic orders. Top right inset describes the proposed criteria for prioritization and the orders identified as conservation and research priorities. Plots show the percentage of species per order for which roadkill data was available in our compilations used as criteria. Order names are followed by the estimated median roadkill rates (rk) calculated from the species rates predicted by trait-based Random Forest models (Medrano-Vizcaíno et al., 2022). Orders are listed from highest to lowest rk with bar color indicating the terciles of the median roadkill rates used as criteria (Brown-orange bars = top tercile, peach bars = medium tercile, and light blue bars = bottom tercile). We also provide for each order the number of species classified by the IUCN as Threatened (T), Data Deficient (DD), Not Threatened (NoT) and the percentage (%) of the total listed as T and DD which was used in the criteria. Animal silhouettes were obtained from Phylopic (http://phylopic.org).

## Methods

### Priority areas

We capitalized on existing data to calculate three metrics for each 1° x 1° grid cell in Latin America (Fig. 1). We defined local vulnerability to road mortality as the sum of predicted roadkill rates for all species occurring in that cell (see details below). Local exposure was calculated as the total length of primary and secondary roads using data from Meijer et al., (2018). Finally, each cell was classified as either studied, if at least one systematic roadkill survey had been conducted in that area (centroid of the study area overlap the cell), or unstudied, if no studies were found. Values for the three metrics in all grid cells are available as supplementary material (Appendices 1-2).

Predicted roadkill rates (number of individual killed per kilometer of road per year, ind/km/year) were available for 346 bird and 159 mammal species reported at least once as roadkill in a recent compilation of systematic road survey studies (Medrano-Vizcaíno et al., 2022). Predicted roadkill rates for each species and 1° x 1° grid cell across Latin America were obtained using trait-based machine learning Random Forest models that related observed roadkill rates from 85 systematic roadkill surveys with study location, species trait data (ecological, life-history, and morphological characteristics) and habitat preferences. Predictions were made for each cell in a species’ area of habitat using current distribution ranges (IUCN, 2021) from which areas without suitable habitat were removed to better represent where species are likely to be present (Brooks et al., 2019). Suitable habitats were defined based on habitat preferences of each species (described in Medrano-Vizcaíno et al., 2022) using high resolution (30 arc-second^2^, ∼1sqkm) land cover data (Latham et al., 2014).

All grid cell values of local vulnerability and local exposure for birds and mammals were grouped into terciles for each metric which combined resulted in nine joint categories (Fig. 1) and were mapped using bivariate choropleth maps in QGIS v.3.18.2-Zürich (https://qgis.org/en/site/). Conservation priority areas were proposed as those falling in the category representing the top tercile of local vulnerability and the bottom tercile of road abundance (low exposure). Top research priorities were defined as unstudied areas that fell in the top terciles of both local vulnerability and local exposure.

### Priority taxa

Using predicted roadkill rates from Medrano-Vizcaíno et al., (2022) and the IUCN Red List categories (IUCN, 2021) we calculated several metrics for each taxonomic order of birds and mammals including: the total number and percentage of species reported as roadkill, the number and percentage of species classified as Threatened (Critically Endangered, Endangered, and Vulnerable), Data Deficient, and Not Threatened (Near Threatened and Least Concern) by the IUCN Red List (IUCN, 2021). We also calculated the average predicted roadkill rate for each order as the median across of all predicted roadkill rates for species in that group. We focused on taxonomic orders as priority targets to illustrate the method, but priorities could be defined at different taxonomic levels or for example considering functional groups. We propose as conservation priorities to orders with at least 1/3 of species recorded as roadkill and high predicted roadkill rates (median roadkill rate in the top tercile of values). We suggest two types of research priorities: type A for understudied but likely susceptible orders (high predicted roadkill rates – median roadkill rate in the top tercile of values, but roadkill data available for <1/3 of species), and type B for unstudied groups of conservation concern (no roadkill data available and at least 1/3 of species listed as Threatened or Data Deficient).

These proposed criteria for priority areas and taxa demonstrate the approach, but different thresholds or criteria could be used depending on the interests, needs, and resources available. To facilitate exploration of priorities under different thresholds for these taxa in Latin America we have made all data available (Appendices 1-4).

## Results

### Priority areas

Local vulnerability and exposure are heterogeneously distributed across Latin America, but vulnerability is generally higher in tropical areas for both birds and mammals (Fig 1). The bivariate maps show priority conservation areas (those with high vulnerability but low exposure) in most of the Amazon region with smaller areas in southern Argentina, northeastern Honduras, and the border of Panama with Colombia (Fig 1). Bird and mammal conservation areas partially overlap but are not identical, reflecting different distributions of species vulnerable to roadkill (Fig 1). Top research priorities (unstudied areas with high vulnerability and high exposure) occurred in most of Central America, northern regions of Venezuela and Colombia, a great part of Ecuador, western Perú, southern Chile, Uruguay, central Argentina, and some coastal areas in Brazil (particularly for birds) (Fig. 1).

At a national scale we note that we found no reported roadkill surveys for five countries that overlap with identified top research priority areas (Belice, El Salvador, Honduras, Nicaragua, and Uruguay).

### Priority taxa

Roadkill estimates were available for only 7.50% and 8.55% of Latin American birds and mammals respectively, and entire orders had no reported roadkill records (Fig 2). From the 29 bird orders present in Latin America, we found data for about two thirds (19) but for represented orders fewer than 50% of the species had reported roadkill estimates. Mammals were better represented with data for species in 10 of the 13 orders in Latin America and data for >50% of species for two orders, but still roadkill estimates reflected a relatively small fraction of mammalian diversity (Fig 2). Median predicted roadkill rates ranged from 0.019 to 0.132 ind/km/year in bird taxonomic orders (SD ranging from 0.018 to 0.129), and from 0.024 to 0.074 in mammalian orders (SD 0.018 to 0.281).

Using our suggested criteria, Cathartiformes and Cariamiformes should be considered as conservation priorities for birds, and Pilosa and Cingulata for mammals (Fig. 2). Cathartiformes are New World vultures that as scavengers can be attracted to roads to forage on roadkill increasing their risk of collision. This group has the highest predicted rate for birds, and more than one third of species (42.86%) have been reported as roadkill. Cathartiformes includes threatened species for which estimates of roadkill are still not available but could be vulnerable to road mortality such as the Andean condor (*Vultur griphus*) and the California condor (*Gymnogyps californianus*). Cariamiformes has the second highest predicted rates for birds, with roadkill data available for one out of two existing species.

Sloths and anteaters (Pilosa) are also found frequently as roadkill (likely their slow movements increase probability of collisions when crossing roads), and more than two thirds of species (80%) have been reported as roadkill. Armadillos (Cingulata) also show vulnerability to roadkill (high rates) and is a group with an uncertain conservation status (five species classified as Data Deficient. Fig. 2).

Cuckoos (Cuculiformes), Caprimulgiformes, Pelecaniformes, and Anseriformes were identified as research priorities type A for birds because roadkill estimates were high, yet risk for many species in the group remains unknown (Fig. 2). Procellariiformes sea birds, and penguins (Sphenisciformes) are research priorities type B because these groups are of conservation concern, but no studies have evaluated roadkill risks. Given these priorities, road ecology research in coastal areas and islands seems particularly necessary to better understand risk for avian species in Latin America. Roadkill rates could be very low for these species, particularly those that spend most of their life at sea but, some species in these groups are known to be affected in other areas. For example, penguins have been reported as roadkill (Heber et al., 2008).

Rabbits (Lagomorpha) were identified as mammalian research priorities type A with high rates combined with a high proportion of unstudied species, while the small mammals in the orders Paucituberculata and Eulipotyphla were considered research priorities type B given the current lack of knowledge and their high proportion of species listed as Threatened and Data Deficient (Fig. 2).

We propose priority taxa for conservation and research at the level of taxonomic orders, however, vulnerability at taxonomic family and species level under different thresholds can be evaluated using data in Appendices 3 and 4 (predicted roadkill rates based on Random Forest models from Medrano-Vizcaíno et al. 2022) available at https://figshare.com/s/5cbce6ffc49953a8866cin.

## Discussion

Our approach capitalizes on existing data to generate quantitative estimates of spatial and taxonomic vulnerability that combined with data on local and taxa-specific risks (e.g., exposure to roads and conservation status) can help identify priorities for conservation and research. We apply this approach to suggest priorities for Latin America based on a set of proposed criteria. These criteria are flexible and could be adapted to reflect different needs, values, and funding trade-offs. Additional sources of risk could also be considered to further optimize conservation and research resources. This approach aims to provide a unified way to predict and map road vulnerabilities, exposure, and risks and facilitate the decision process of where and who to protect and study at a large scale. We envision this as a first step in the decision-making process that could complement existing conservation prioritization tools (Schwartz et al., 2020; Zizka et al., 2021) and should be followed with local and regional analyses to develop conservation and research agendas.

Application of the proposed approach to define conservation priorities in Latin America revealed the Amazon as a focal region to minimize development of infrastructure. The presence of many species with high predicted roadkill rates means that expanding the road network in this area without careful planning and adequate mitigation measures will likely have negative consequences for both wildlife and humans, as collisions can result in injuries and even fatalities (Zhao et al., 2010). The Amazon is already considered a priority for conservation due to its unique forest environment, carbon sequestering potential and high biodiversity (Strassburg et al., 2010). New roads would likely lead to increased wildlife mortality, but even if mitigation measures were implemented to reduce mortality, new roads facilitate further degradation and human expansion (Laurance et al., 2009). Worryingly, 12,263 km of roads may be opened or improved (leading to more traffic and higher travelling speeds) in the Amazon in the next few years, and this expansion will likely be accompanied by additional illegal roads which can triple the length of official roads in some areas (Barber et al., 2014; Vilela et al., 2020).

Applying the proposed approach to Latin America also suggested priority areas for future studies, particularly in Central America and North-western South America. Research efforts can be driven by individual interests, but funding agencies can also define priorities and propose targeted schemes. We hope that by identifying research priorities in Latin America, where expertise is expanding but data are still limited compared to other regions (Silva et al., 2021), our approach will encourage researchers and funding agencies to focus on understudied areas where species are more susceptible to road impacts and roads abound. These are areas where knowledge is most urgently needed to quantify risks and if necessary, propose mitigation measures.

Our analyses also identified taxonomic groups as conservation or research priorities. Further evaluation of species in these target groups may reveal limited risk for some due to behavioral avoidance [which can prevent roadkill but still have negative impacts by reducing gene flow (Holderegger & Di Giulio, 2010)] or preferences for local habitats where roads are absent or rare and thus, risks are few. Deeper evaluation of the proposed, admittedly coarse, priorities would help refine these targets, eliminating taxa unlikely to encounter roads or currently limited to roadless areas. We recognize that targeting taxa during roadkill assessments could present some challenges, as some species are naturally more difficult to detect even if roadkilled. For example, species-level roadkill data for smaller mammals like rodents or shrews is rare, likely because small animals are more often unrecognizable after collision – found as hairy spots on roads if at all (Cook & Blumstein, 2013). Additionally, rarer groups of small mammals (e.g., Eulipotyphla and Paucituberculata) may be wrongly classified in broad groups (like “unclassified rodents”) which prevents species-specific analyses. Future research should explore ways to address these biases. Improving taxonomic identification skills or consulting with experts can be a solution. Although expensive, another solution would be identification of roadkill samples using molecular techniques.

The Latin America and Caribbean region is home to more than 4600 birds and 1800 mammals (IUCN, 2021), yet estimates of mortality caused by collisions with vehicles are available for <10% of species (Medrano-Vizcaíno et al., 2022). This means that the magnitude of how roads impact wildlife is likely underestimated. Finding ways to identify priority areas and taxa for research can address this limitation more efficiently to improve knowledge. At the same time, roadkill surveys can provide information about rare and poorly studied species. For example, the western mountain coati *Nasuella olivacea*, considered the least-studied carnivore of the world (Helgen et al., 2009), was the second most roadkilled species in a study in Colombia (Delgado-V, 2007). Specimens collected as roadkill can contribute to understand the biology of organisms that can be difficult to study, making roadkill assessment valuable beyond the estimate of threats (Medrano-Vizcaíno & Brito-Zapata, 2021).

Latin America is not the only region where future road development and lack of road impact assessment coincide. In Asia, systematic studies focused on wildlife mortality have been conducted in only nine out of 48 countries, while in Africa there is systematic data for only five out of 54 countries (Silva et al., 2021), yet rapid development of infrastructure is expected in these regions (Meijer et al., 2018). While funds to assess potential risk may not always accompany road development plans, identifying regions where this research is particularly needed could aid in more directed investment of limited resources. Our approach could be applied to different taxonomic or functional groups and areas of the world to better plan research and conservation actions. Our study reveals priority areas and taxa that we hope would be valuable to ecologists, funding agencies, and decision makers in Latin America. Additionally, we provide a comprehensive dataset of predicted mortality rates of birds and mammals across all Latin America, which can be used to propose priorities for different taxonomic levels and geographic scales in this region.

## Acknowledgements

We thank University of Reading for funding this work through an International Research Studentship for PhD awarded to PMV in 2019 (ref GS19-042). We also acknowledge financial support to CESAM by FCT/MCTES (UIDP/50017/2020+UIDB/50017/2020+ LA/P/0094/2020), through national funds.

## Data availability statement

All data used for analyses including birds and mammals’ vulnerability to roadkills and the grid for Latin America are available as appendices at: https://figshare.com/s/5cbce6ffc49953a8866c.

## References

Barber, C. P., Cochrane, M. A., Souza, C. M., & Laurance, W. F. (2014). Roads, deforestation, and the mitigating effect of protected areas in the Amazon. Biological Conservation, 177(2014), 203–209. https://doi.org/10.1016/j.biocon.2014.07.004

Brooks, T. M., Pimm, S. L., Resit Akç, H., Buchanan, G. M., Butchart, S. H. M., Foden, W., Hilton-Taylor, C., Hoffmann, M., Jenkins, C. N., Joppa, L., Li, B. V, Menon, V., Ocampo-Peñ, N., & Rondinini, C. (2019). Measuring Terrestrial Area of Habitat (AOH) and Its Utility for the IUCN Red List The International Union for Conservation of Nature (IUCN) Red List of Threatened Species includes assessment of extinction risk for. Trends in Ecology & Evolution, 34(11). https://doi.org/10.1016/j.tree.2019.06.009

Cook, T. C., & Blumstein, D. T. (2013). The omnivore’s dilemma: Diet explains variation in vulnerability to vehicle collision mortality. Biological Conservation, 167, 310–315. https://doi.org/10.1016/j.biocon.2013.08.016

Delgado-V, C. (2007). Muerte de mamíferos por vehículos en la vía del Escobero, Envigado (Antioquia), Colombia. Actual Biol, 29(87), 235–239.

González-Suárez, M., Zanchetta Ferreira, F., & Grilo, C. (2018). Spatial and species-level predictions of road mortality risk using trait data. Global Ecology and Biogeography, 27(9), 1093–1105. https://doi.org/10.1111/geb.12769

Grilo, C., Koroleva, E., Andrášik, R., Bíl, M., & González-Suárez, M. (2020). Roadkill risk and population vulnerability in European birds and mammals. Frontiers in Ecology and the Environment, 18(6), 323–328. https://doi.org/10.1002/fee.2216

Heber, S., Wilson, K. J., & Molles, L. (2008). Breeding biology and breeding success of the blue penguin (Eudyptula minor) on the West Coast of New Zealand’s South Island. New Zealand Journal of Zoology, 35(1), 63–71. https://doi.org/10.1080/03014220809510103

Helgen, K. M., Kays, R., Helgen, L. E., Tsuchiya-Jerep, M. T. N., Pinto, C. M., Koepfli, K.-P., Eizirik, E., & Maldonado, J. E. (2009). Taxonomic boundaries and geographic distributions revealed by an integrative systematic overview of the mountain coatis, Nasuella (Carnivora: Procyonidae). Small Carnivore Conservation, 41, 65–74.

Hill, J. E., DeVault, T. L., & Belant, J. L. (2019). Cause-specific mortality of the world’s terrestrial vertebrates. Global Ecology and Biogeography, 28(5), 680–689. https://doi.org/10.1111/geb.12881

Holderegger, R., & Di Giulio, M. (2010). The genetic effects of roads: A review of empirical evidence. Basic and Applied Ecology, 11(6), 522–531. https://doi.org/10.1016/j.baae.2010.06.006

IUCN. (2021). The IUCN Red List of Threatened Species. Version 2021-1. https://www.iucnredlist.org

Laurance, W. F., Clements, G. R., Sloan, S., O’Connell, C. S., Mueller, N. D., Goosem, M., Venter, O., Edwards, D. P., Phalan, B., Balmford, A., Van Der Ree, R., & Arrea, I. B. (2014). A global strategy for road building. Nature, 513(7517), 229–232. https://doi.org/10.1038/nature13717

Laurance, W. F., Goosem, M., & Laurance, S. G. W. (2009). Impacts of roads and linear clearings on tropical forests. Trends in Ecology and Evolution, 24(12), 659–669. https://doi.org/10.1016/j.tree.2009.06.009

Loss, S. R., Will, T., & Marra, P. P. (2014). Estimation of bird-vehicle collision mortality on U.S. roads. The Journal of Wildlife Management, 78(5), 763–771. https://doi.org/10.1002/JWMG.721

Medrano-Vizcaíno, P., & Brito-Zapata, D. (2021). Filling biogeographical gaps through wildlife roadkills: New distribution records for six snake species from Ecuador (Anilius scytale, Drymarchon corais, Erythrolamprus breviceps, Micrurus lemniscatus, Oxyrhopus vanidicus, Trilepida anthracina). Neotropical Biodiversity, 7(1), 554–559. https://doi.org/10.1080/23766808.2021.2010469

Medrano-Vizcaíno, P., Grilo, C., Silva, F. A., Carvalho, W. D., Melinski, R. D., & González-Suárez, M. (2022). Roadkill patterns in Latin American birds and mammals. Global Ecology and Biogeography, 0, 1–28. https://doi.org/10.1111/geb.13557

Meijer, J. R., Huijbregts, M. A. J., Schotten, K. C. G. J., & Schipper, A. M. (2018). Global patterns of current and future road infrastructure. Environmental Research Letters, 13(6). https://doi.org/10.1088/1748-9326/aabd42

Myers, N., Mittermeler, R. A., Mittermeler, C. G., Da Fonseca, G. A. B., & Kent, J. (2000). Biodiversity hotspots for conservation priorities. Nature, 403(6772), 853–858. https://doi.org/10.1038/35002501

Pinto, F. A. S., Clevenger, A. P., & Grilo, C. (2020). Effects of roads on terrestrial vertebrate species in Latin America. Environmental Impact Assessment Review, 81, 106337. https://doi.org/10.1016/j.eiar.2019.106337

Schwartz, A. L. W., Shilling, F. M., & Perkins, S. E. (2020). The value of monitoring wildlife roadkill. European Journal of Wildlife Research, 66(1), 1–12. https://doi.org/10.1007/s10344-019-1357-4

Silva, I., Crane, M., & Savini, T. (2021). The road less traveled: Addressing reproducibility and conservation priorities of wildlife-vehicle collision studies in tropical and subtropical regions. Global Ecology and Conservation, 27, 1–12. https://doi.org/10.1016/j.gecco.2021.e01584

Strassburg, B. B. N., Kelly, A., Balmford, A., Davies, R. G., Gibbs, H. K., Lovett, A., Miles, L., Orme, C. D. L., Price, J., Turner, R. K., & Rodrigues, A. S. L. (2010). Global congruence of carbon storage and biodiversity in terrestrial ecosystems. Conservation Letters, 3, 98–105. https://doi.org/10.1111/j.1755-263X.2009.00092.x

Taylor-Brown, A., Booth, R., Gillett, A., Mealy, E., Ogbourne, S. M., Polkinghorne, A., & Conroy, G. C. (2019). The impact of human activities on Australian wildlife. PLoS ONE, 14(1), 1–28. https://doi.org/10.1371/journal.pone.0206958

Vilela, T., Harb, A. M., Bruner, A., Da Silva Arruda, V. L., Ribeiro, V., Alencar, A. A. C., Grandez, A. J. E., Rojas, A., Laina, A., & Botero, R. (2020). A better Amazon road network for people and the environment. Proceedings of the National Academy of Sciences of the United States of America, 117(13), 7095–7102. https://doi.org/10.1073/pnas.1910853117

Zhao, H., Yin, Z., Chen, R., Chen, H., Song, C., Yang, G., & Wang, Z. (2010). Investigation of 184 passenger car-pedestrian accidents. International Journal of Crashworthiness, 15(3), 313–320. https://doi.org/10.1080/13588260903335290

Zizka, A., Barratt, C. D., Ritter, C. D., Joerger-Hickfang, T., & Zizka, V. M. A. (2021). Existing approaches and future directions to link macroecology, macroevolution and conservation prioritization. Ecography, 44, 1–15. https://doi.org/10.1111/ecog.05557

